# Molecular basis for the recognition of the human AAUAAA polyadenylation signal

**DOI:** 10.1101/209809

**Authors:** Yadong Sun, Yixiao Zhang, Keith Hamilton, James L. Manley, Yongsheng Shi, Thomas Walz, Liang Tong

## Abstract

Nearly all eukaryotic messenger RNA precursors must undergo cleavage and polyadenylation at their 3′-end for maturation. A crucial step in this process is the recognition of the AAUAAA polyadenylation signal (PAS), and the molecular mechanism of this recognition has been a long-standing problem. Here we report the cryo-electron microscopy structure of a quaternary complex of human CPSF-160, WDR33, CPSF-30 and an AAUAAA RNA at 3.4 Å resolution. Strikingly, the AAUAAA PAS assumes an unusual conformation that allows this short motif to be bound directly by both CPSF-30 and WDR33. The A1 and A2 bases are recognized specifically by zinc finger 2 (ZF2) of CPSF-30 and the A4 and A5 bases by ZF3. Interestingly, the U3 and A6 bases form an intramolecular Hoogsteen base pair and directly contact WDR33. CPSF-160 functions as an essential scaffold and pre-organizes CPSF-30 and WDR33 for high-affinity binding to AAUAAA. Our findings provide an elegant molecular explanation for how PAS sequences are recognized for mRNA 3′-end formation.

## Introduction

In eukaryotes, messenger RNAs are transcribed from the genome by RNA polymerase II (Pol II). However, these primary transcripts (pre-mRNAs) must undergo extensive processing, including 5′-end capping and splicing. The pre-mRNAs must also be processed at their 3′ end. For most of them, the processing involves an endonucleolytic cleavage at a specific location followed by the addition of a poly(A) tail (polyadenylation) (1–5). Metazoan replication-dependent histone pre-mRNAs are distinct in that they are only cleaved but not polyadenylated (6, 7). The mature mRNAs are then transported into the cytoplasm, where they can be translated into proteins.

A large number of protein factors have been identified for pre-mRNA 3′-end processing (8–10). The machinery for canonical 3′-end processing (cleavage and polyadenylation) has been fractionated into several components based on classical biochemistry experiments, including the cleavage and polyadenylation specificity factor (CPSF) (11, 12) and the cleavage stimulation factor (CstF) (13, 14) in mammals. CPSF has six subunits and has central roles in 3′-end processing. Its CPSF-73 subunit is the endoribonuclease for the cleavage reaction (15), and its WDR33 and CPSF-30 subunits are required for recognizing the polyadenylation signal (PAS) in the pre-mRNA (16, 17), which helps define the position of RNA cleavage. The PAS consists of a hexanucleotide, most frequently with the sequence AAUAAA (18, 19) and first identified more than 40 years ago (20), and is typically located 10-30 nucleotides upstream of the cleavage site. CstF has three subunits and recognizes a G/U-rich sequence motif downstream of the cleavage site. It also contributes to the selection of the cleavage site, including alternative polyadenylation (21-23).

The subunits of CPSF form two functional components. mPSF (mammalian polyadenylation specificity factor) consists of CPSF-160, WDR33, CPSF-30 and Fip1, and is necessary and sufficient for PAS recognition and polyadenylation (with poly(A) polymerase) (17). CPSF-73 and CPSF-100 together with symplekin form the other component, also known as the core cleavage complex (24) or mCF (mammalian cleavage factor) (5), which catalyzes the cleavage reaction. Symplekin is a scaffold protein and also mediates interactions with CstF and other factors in the 3′-end processing machinery (25–28).

CPSF-160 (160 kDa) contains three β-propeller domains (BPA, BPB and BPC) and a C-terminal domain (CTD) (Fig. 1A), and shares weak sequence homology to the DNA damage-binding protein (DDB1) (29, 30). WDR33 (146 kDa) contains a WD40 domain near the N terminus, and a collagen-like segment near the middle (10). CPSF-30 (30 kDa) contains five CCCH zinc-finger motifs (ZF1-ZF5) near the N terminus and a CCHC zinc knuckle at the C terminus (31).

**Figure 1.**
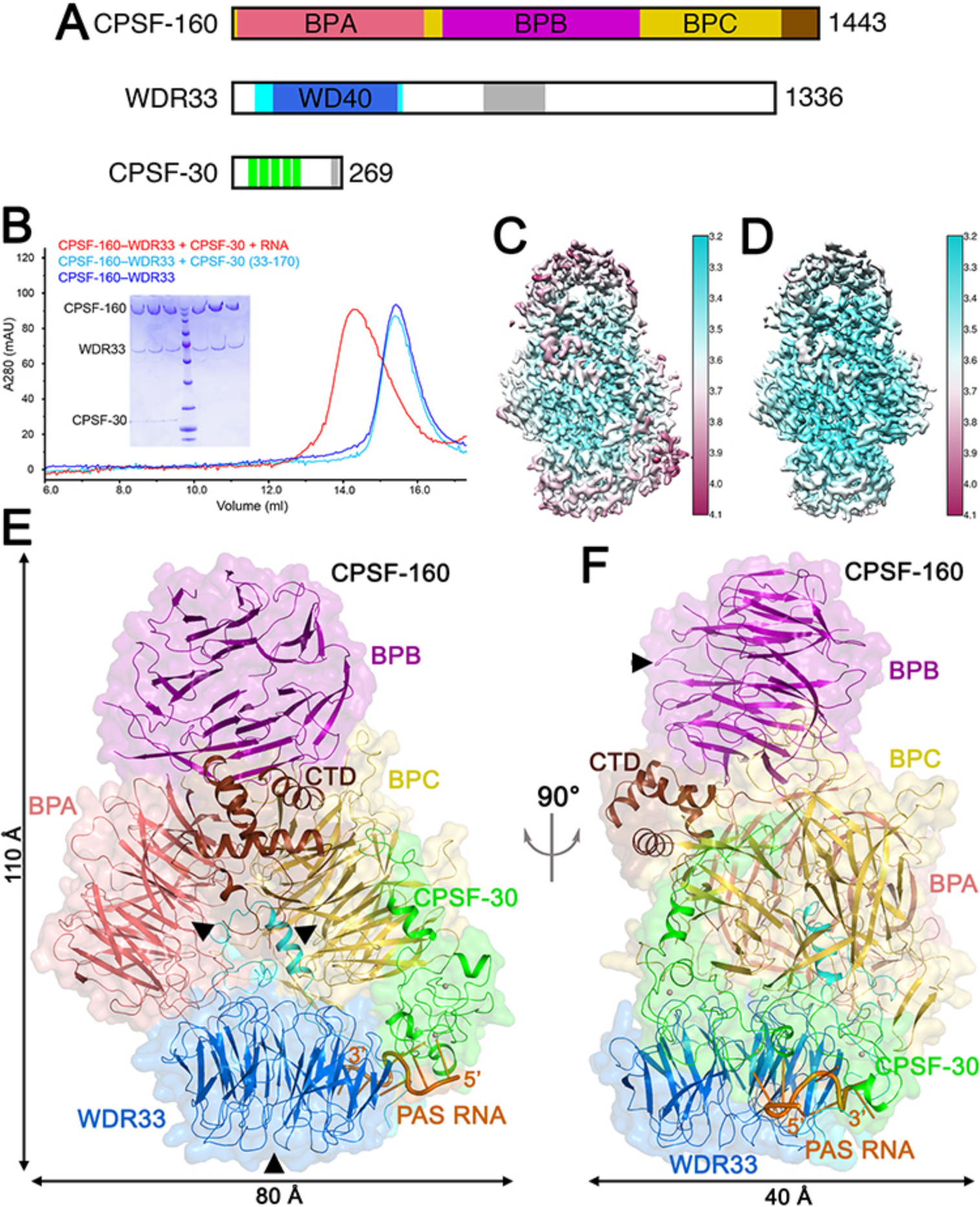
Overall structure of the human CPSF-160–WDR33–CPSF-30–PAS RNA quaternary complex. **(A)**. Domain organizations of human CPSF-160, WDR33 and CPSF-30. The β-propeller domains of CPSF-160 are labeled and given different colors, and the CTD in brown. The WD40 domain of WDR33 is in light blue, and the surrounding segments in cyan. The collagen-like segment is in gray. ZF1-ZF5 in CPSF-30 are in green, and the zinc knuckle in gray. **(B)**. Gel filtration profiles of the quaternary complex (red), the CPSF-160–WDR33 binary complex (blue), and a mixture of CPSF-160, WDR33, and a CPSF-30 deletion mutant (light blue). Inset: SDS-PAGE gel of the quaternary and binary complex samples. **(C)**. Local resolution map for the CPSF-160–WDR33–CPSF-30–PAS RNA quaternary complex. **(D)**. Local resolution map for the CPSF-160–WDR33 binary complex. **(E)**. Overall structure of the quaternary complex, colored as in panel A. The PAS RNA is in orange. The molecular surface of the structure is shown as a transparent surface. The top faces of BPA and BPC of CPSF-160 and the WD40 domain of WDR33 are indicated with black arrowheads. **(F)**. Overall structure of the quaternary complex, viewed after a rotation of 90° around the vertical axis. Panels C and D were produced with Chimera (66), and the other structure figures were produced with PyMOL (http://www.pymol.org).

The recognition of the PAS is a critical step in canonical 3′-end processing. However, its molecular mechanism is not known, and this has been a long-standing question in the field. Here we have determined the structure of a quaternary complex of human WDR33, CPSF-30, CPSF-160 and a PAS RNA by cryogenic electron microscopy (cryo-EM) at 3.4 Å resolution. Most importantly, the structure has illuminated the molecular basis for the specific recognition of the PAS, revealing direct roles of CPSF-30 and WDR33, as well as extensive interactions among the proteins factors in this complex, which pre-organizes the binding site for the PAS RNA.

## Results

### Structure determination

We produced the complex of full-length human CPSF-160 with the N-terminal region of human WDR33 (residues 1-572) by co-expressing them in baculovirus-infected insect cells (Fig. 1B) and studied the sample by EM. We produced full-length human CPSF-30 fused to maltose-binding protein (MBP) by expression in bacteria. To prepare the quaternary complex, the three proteins were mixed together with a 17-mer RNA oligo containing a PAS (5′-AACCUCC<u>AAUAAA</u>CAAC-3′, with the PAS underlined), incubated with a protease to remove the MBP, and the complex was purified by gel filtration chromatography for EM (Fig. 1B).

We determined the structures of the CPSF-160–WDR33 binary complex at 3.8 Å resolution by cryo-EM (Figs. S1-S3), which allowed us to build an atomic model of the complex. We next determined the structure of the CPSF-160–WDR33–CPSF-30–PAS RNA quaternary complex at 3.4 Å resolution (Fig. 1C, Figs. S2, S3). Compared to CPSF-160 and WDR33, the resolution of the CPSF-30 and RNA region is somewhat lower, suggesting that these two molecules may have some overall flexibility in the quaternary complex. We also determined a higher quality structure of the CPSF-160–WDR33 binary complex, at 3.3 Å resolution, based on the cryo-EM images from the quaternary complex sample (Fig. 1D, Figs. S2, S3). The atomic models for the three structures have good fit to the EM density as well as expected geometric parameters (Table S1, Fig. S3).

### Overall structure of the quaternary complex

The overall structure of the quaternary complex has dimensions of 110 Å × 80 Å × 40 Å (Figs. 1E, 1F). The three β-propellers of CPSF-160 are arranged in a trefoil pattern, and its CTD is located near the center of the trefoil, interacting with the sides of the three propellers. WDR33 is bound between the ‘top’ faces of BPA and BPC. The top face is where the loops connecting neighboring blades of a propeller are located (30). CPSF-30 is located on one side of the structure, contacting both BPC in CPSF-160 and WDR33. The PAS RNA is positioned between CPSF-30 and WDR33, and has no direct interactions with CPSF-160.

Twelve surface segments of CPSF-160 (containing 266 of the 1,443 residues) are not included in the atomic model of the quaternary complex due to lack of density, suggesting that they are likely disordered. Residues 1-42 and 418-572 of WDR33 are disordered in the quaternary complex. Residues 117-269 of CPSF-30 are also disordered, which include ZF4, ZF5 and the zinc knuckle. These residues of CPSF-30 may mediate other functions in the pre-mRNA 3′-end processing machinery, although the yeast homolog Yth1 does not contain a zinc knuckle.

### Interaction between CPSF-160 and WDR33

CPSF-160 and WDR33 have an extensive interface, burying approximately 3,700 Å^2^ of the surface area of each protein (Fig. 2A). For WDR33, the interaction with CPSF-160 is primarily mediated by the segment before its WD40 domain, residues 52-109, which contribute 2,500 Å^2^ to the buried surface area. These residues are located in a deep pocket at the BPA-BPC interface of CPSF-160 (Figs. 1E, 2B). The loops on the bottom face of the WD40 domain contribute 850 Å^2^ to the buried surface area, and they contact loops on the top faces of BPA and BPC (Fig. 1E). The segment C-terminal to the WD40 domain (residues 403-417) contributes 250 Å^2^, and these residues are located near the segment before the WD40 domain (Fig. 2B). Residues 41-109 and 403-420 are well conserved, while residues 1-40 are poorly conserved among WDR33 homologs (Fig. S4).

**Figure 2.**
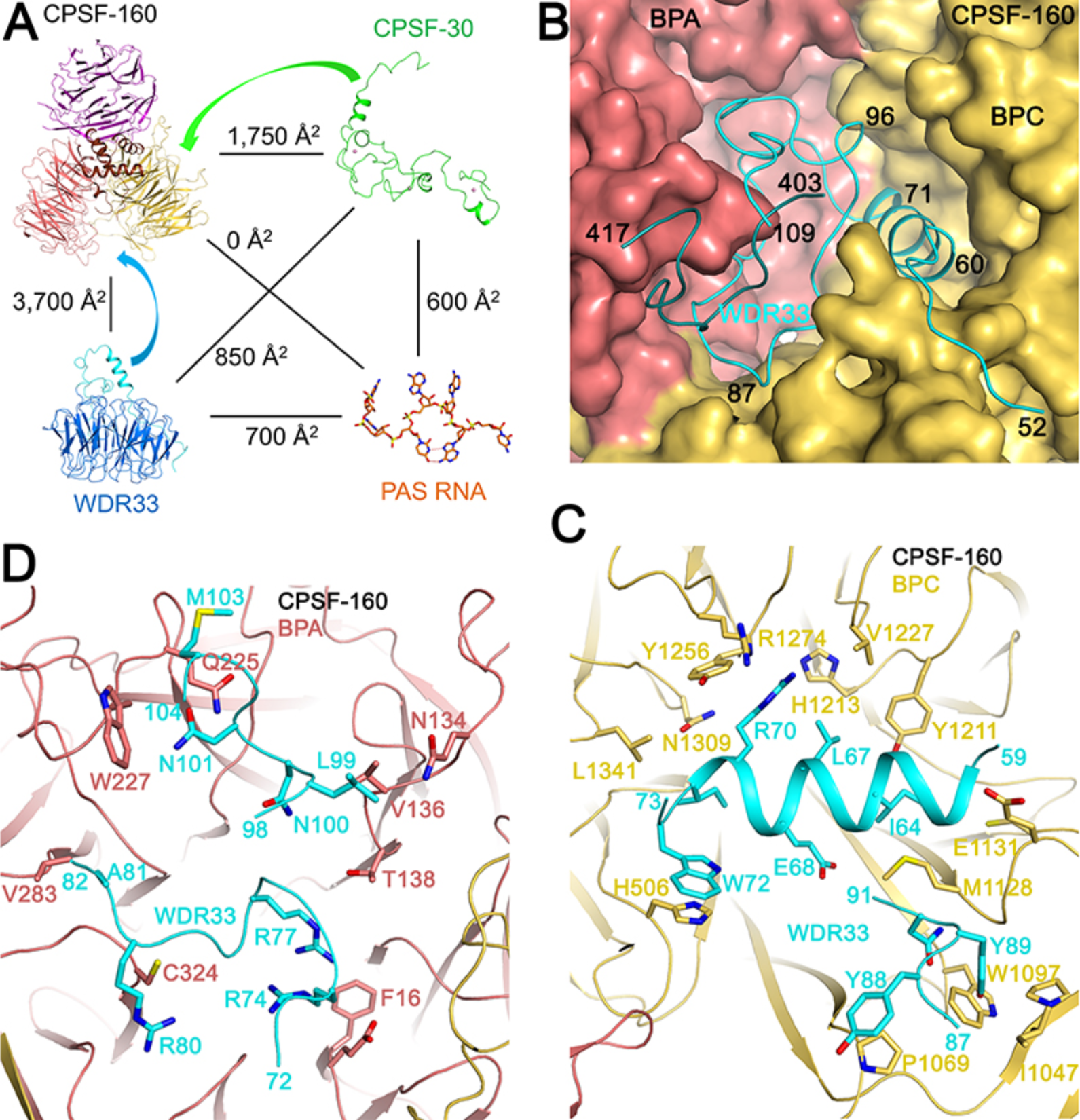
Interactions between human CPSF-160 and WDR33. **(A)**. Buried surface areas at the interfaces in the CPSF-160–WDR33–CPSF-30–PAS RNA quaternary complex. Curved arrows indicate the major contacts at the CPSF-160–WDR33 and CPSF-160– CPSF-30 interfaces. **(B)**. Molecular surface of CPSF-160, colored by domains. The N-and C-terminal segments of WDR33 beyond the WD40 domain, shown as cartoons in cyan and dark cyan, respectively, are located in a deep pocket in CPSF-160. **(C)**. Interactions between WDR33 (residues 59-73 and 87-91, cyan) with the top face of BPC of CPSF-160 (yellow). Most of the residues shown in sticks contribute >50 Å^2^ to the buried surface area at the interface. **(D)**. Interactions between WDR33 (residues 72-82 and 98-104, cyan) with the top face of BPA of CPSF-160 (salmon).

The interaction between the N-terminal segment of WDR33 and BPC involves residues 52-73 and 85-91 of WDR33 and is more extensive than that with BPA of CPSF-160, which involves residues 74-84 and 99-104 of WDR33. Especially, residues 60-71 of WDR33 form a helix and are positioned near the center of the BPC top face, showing hydrophobic as well as polar interactions (Fig. 2C). The loop containing residues 87-91 has intimate contacts with two blades of BPC, at one side of the top face (Fig. 2C). In contrast, a loop containing residues 72-82 of WDR33 is located near the center of the BPA top face, with two Arg residues (Arg74 and Arg77) projecting into the central cavity of the propeller (Fig. 2D). However, these two residues do not establish strong interactions with BPA, as they do not have clearly favorable binding partners in CPSF-160. The loop containing residues 98-104 of WDR33 is positioned at one side of the top face of BPA, interacting with two blades of the propeller (Fig. 2D).

The large surface area buried at the CPSF-160–WDR33 interface suggests that their binary complex should be stable as well. In fact, the overall structure of the CPSF-160– WDR33 binary complex is essentially the same as that in the quaternary complex (Fig. S5), with an rms distance of 0.24 Å for 1,539 equivalent Cα atoms. Therefore, binding of CPSF-30 and PAS RNA does not cause a global structural change in the CPSF-160–WDR33 complex, except that residues 43-54 of WDR33 are disordered in the binary complex. They become ordered in the quaternary complex and contact the PAS RNA and ZF3 of CPSF-30 (Fig. S5; see below).

### Interaction of CPSF-30 with CPSF-160 and WDR33

The interface between CPSF-160 and CPSF-30 buries 1,700 Å^2^ of the surface area of each protein (Fig. 2A). Residues 1-26 of CPSF-30, in an N-terminal extension before the zinc fingers (Fig. 1A) and well conserved among homologs (Fig. S6), contact the bottom face of BPC (Figs. 1E, 3A), contributing 1,100 Å^2^ to the buried surface area, indicating that these residues are crucial for the interactions between CPSF-160 and CPSF-30. Especially, residues 1-7 of CPSF-30 are located in a deep pocket at the center of the BPC bottom face (Fig. 3B). Consistent with the structural observations, we found that a CPSF-30 mutant missing the first 32 residues (as well as C-terminal residues disordered in the quaternary complex structure) could not form a ternary complex with CPSF-160 and WDR33 (Fig. 1B).

**Figure 3.**
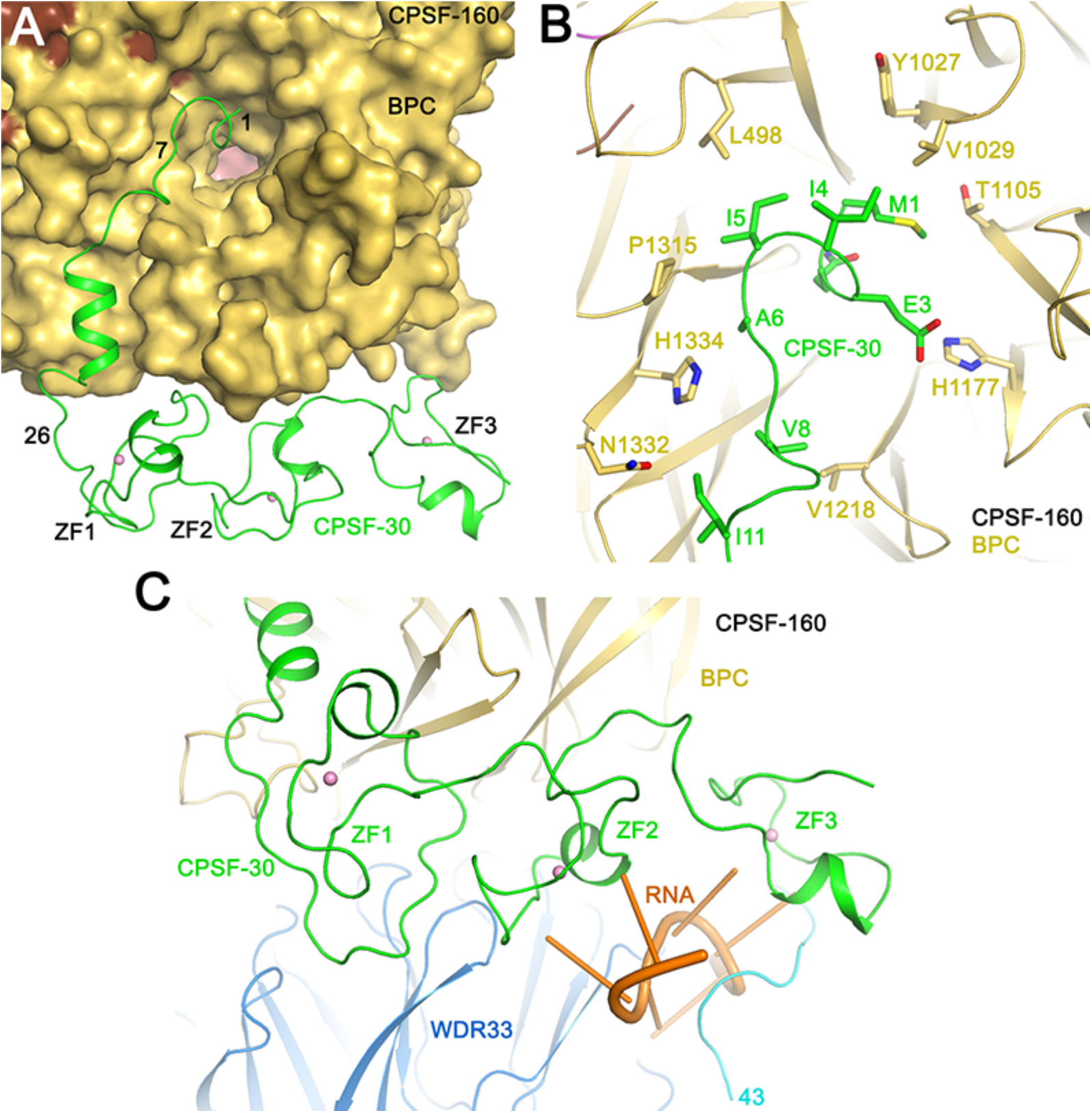
Interactions of human CPSF-30 with CPSF-160 and WDR33. **(A)**. Molecular surface of CPSF-160, colored by domains. CPSF-30 is shown as a cartoon in green, and the metal ions in the zinc fingers as spheres (pink). **(B)**. Interactions between CPSF-30 (residues 1-11, green) with the bottom face of BPC of CPSF-160 (yellow). Most of the residues shown in sticks contribute >40 Å^2^ to the buried surface area at the interface. **(C)**. ZF1-ZF3 of CPSF-30 (green) contact the side of BPC of CPSF-160 (yellow) and the top face of the WDR33 WD40 domain (light blue). The PAS RNA is in orange.

ZF1-ZF3 of CPSF-30 contact both WDR33 (850 Å^2^ buried surface area) and the BPC of CPSF-160 (600 Å^2^, Fig. 3C). The smaller buried surface area here suggests that the interactions are somewhat weaker, although the PAS RNA is situated between WDR33 and CPSF-30 and is likely to stabilize their interactions. ZF1-ZF2 contacts the side of BPC and the edge of the bottom face of the WDR33 WD40 domain, while ZF3 primarily interacts with the N-terminal segment of WDR33 (residues 43-54) that becomes ordered in the quaternary complex (Fig. 3C).

### Binding mode of PAS RNA

While a 17-mer RNA oligo was used for making the quaternary complex, the cryo-EM map revealed density for only 7 nucleotides, the entire AAUAAA PAS and the following nucleotide, which has weak density (Fig. 4A). The fact that only the AAUAAA of the RNA oligo is well ordered in the structure is consistent with this complex being responsible for recognizing the PAS for pre-mRNA 3′-end processing. The nucleotide following the PAS is mostly exposed to the solvent, and will not be described further here.

**Figure 4.**
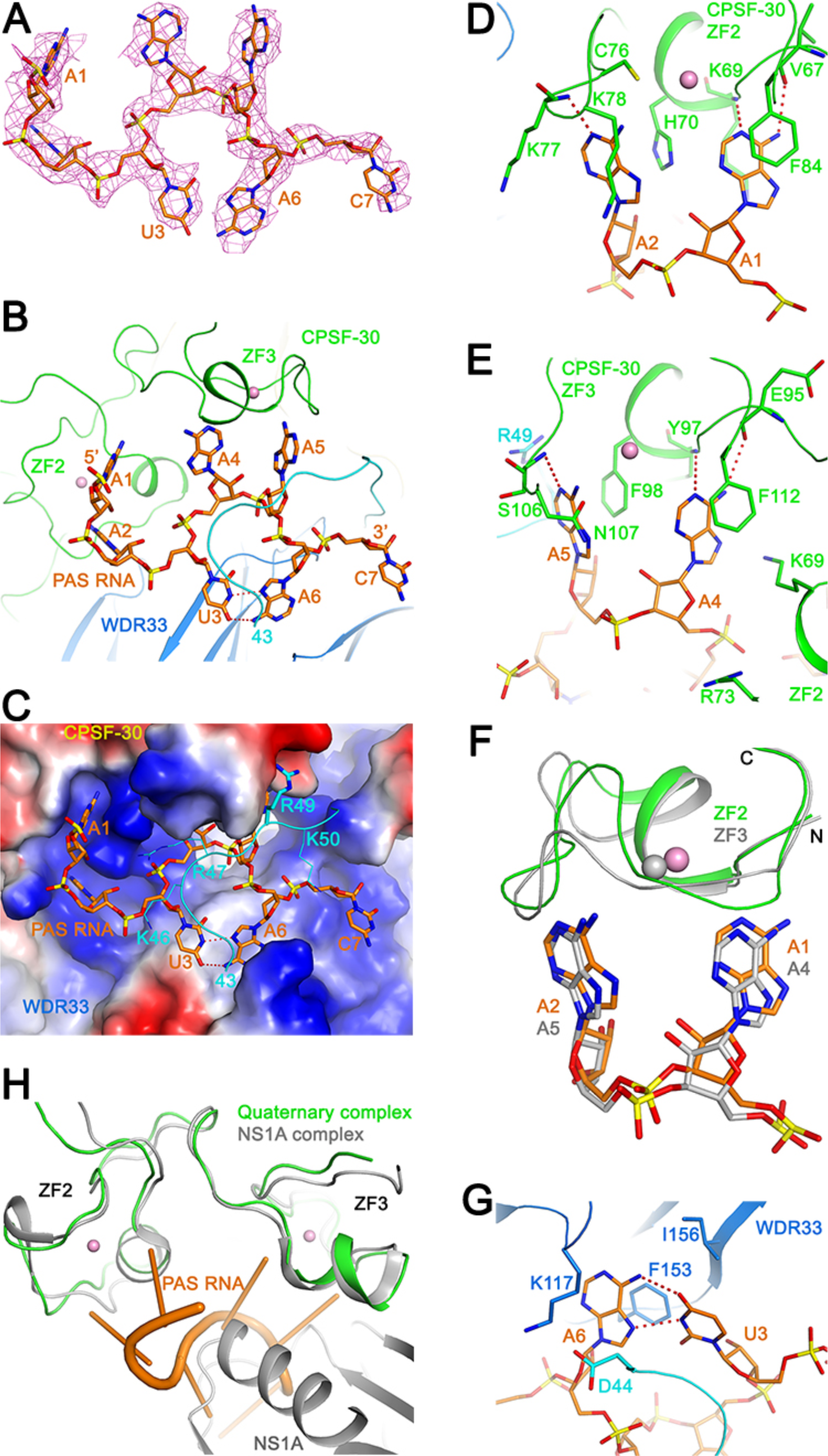
Recognition of the PAS RNA by CPSF-30 and WDR33. **(A)**. Cryo-EM density (magenta mesh) for residues AAUAAAC of the PAS RNA (orange) used in this study, contoured at 4.5σ. **(B)**. Binding mode of the AAUAAA PAS (orange) at the interface of CPSF-30 ZF2-ZF3 (green) and WDR33 (light blue for WD40 domain and cyan for N-terminal segment). Hydrogen-bonds in the U3-A6 Hoogsteen base pair are indicated with dashed lines in red. **(C)**. Molecular surface of CPSF-160–WDR33–CPSF-30 complex, colored by electrostatic potential (red: negative, blue: positive). The N-terminal segment of WDR33 (residues 43-52) is shown as a cartoon only. The side chains of Lys46, Arg47 and Lys50 have no density and are shown as lines. The side chain of Arg49 has density and is shown as sticks. PAS RNA is shown as stick model in orange. **(D)**. Detailed interactions showing the recognition of the A1 and A2 bases of the PAS (orange) by ZF2 of CPSF-30 (green). Hydrogen-bonds between the adenine bases and the protein are indicated with dashed lines in red. **(E)**. Detailed interactions showing the recognition of the A4 and A5 bases of the PAS (orange) by ZF3 of CPSF-30 (green). **(F)**. Overlay of the structures of ZF2 (green) and ZF3 (gray) of CPSF-30 brings their cognate dinucleotides (orange and gray, respectively) into overlap as well. **(G)**. Recognition of the U3-A6 base pair (orange) by WDR33 (blue). **(H)**. Overlay of the structure of ZF2-ZF3 of CPSF-30 in the quaternary complex (green) with that in the complex with influenza virus NS1A protein (PDB entry 2RHK) (41). NS1A is shown in gray, clashing with the PAS RNA.

The AAUAAA PAS is located between the side of the WD40 domain of WDR33 and ZF2-ZF3 of CPSF-30 (Figs. 1E, 4B). The six bases are positioned in clearly defined pockets, while the backbone phosphates are mostly exposed to the solvent, although there are many positively charged residues near the RNA (Fig. 4C). Residues 43-54 in the N-terminal region of WDR33 become ordered in the quaternary complex, and cover up part of the RNA, although the density for this segment is relatively weak (Fig. 4C).

The backbone of the AAUAAA PAS assumes an S shape, and there are no stacking interactions among the six bases, which are all in the *anti* conformation (Fig. 4B). Interestingly, we found that U3 and A6 form a Hoogsteen U-A base pair (the nucleotides in the PAS are numbered from 1 to 6 here). The six bases in the PAS are arranged such that A1 and A2 are pointed in one direction, and are recognized by ZF2 of CPSF-30 (Fig. 4B). A4 and A5 are pointed in another direction, perpendicular to that of A1-A2, and these two bases are recognized by ZF3 of CPSF-30. The U3-A6 base pair is pointed in the opposite direction from that of A4-A5, and is bound by WDR33. With this arrangement, the bases of the trinucleotide A2-U3-A4 are splayed apart, and such conformations have been observed in other protein-RNA complexes (32-34).

### Specific recognition of PAS RNA

The structure reveals that A1, A2, A4 and A5 of the PAS are recognized specifically by CPSF-30. The N1 and N6 atoms of the A1 base are recognized by hydrogen-bonds with the main-chain amide of Lys69 and main-chain carbonyl of Val67 in ZF2, respectively (Fig. 4D), a hydrogen-bonding pattern that is identical to that in an A-U Watson-Crick base pair as well as specific recognition of adenine base by other proteins. The N1 atom of the A2 base is recognized by a hydrogen-bond with the main-chain amide of Lys77, while its N6 atom does not appear to be recognized. In addition, the A1 base is π-stacked with the side chain of Phe84 on one face and flanked by that of Lys69 on the other (Fig. 4D). The A2 base is π-stacked with the side chain of His70 on one face and surrounded by Lys77 and Lys78 on the other. Both bases are also located near the metal ion and its ligands in ZF2.

Remarkably, the A4 and A5 bases are recognized by ZF3 of CPSF-30 using similar interactions (Fig. 4E). In fact, the two zinc fingers show a conserved mechanism of recognizing the bases. An overlay of ZF3 with ZF2 also brings their cognate bases into overlap (Fig. 4F). The geometry of these hydrogen-bonds is not optimal, possibly due in part to the limited resolution of the current structure. It might also be possible that some of these sub-optimal interactions are genuine structural features, which would allow some variability in the nucleotides in the PAS. Nonetheless, the hydrogen-bonding pattern clearly indicates that adenines would be preferred at these positions.

The U3-A6 base pair is π-stacked with the side chain of Phe153 in the WD40 domain of WDR33, while its other face is in contact with residues 43-45 in its N-terminal segment (Fig. 4G). Lys117 and Ile156 surround the side of this base pair. While these two bases do not appear to be specifically recognized directly, a G6 base would not be preferred here as its 2-amino group would clash with the main-chain carbonyl of Thr115. Residues that are in contact with the RNA are highly conserved among WDR33 and CPSF-30 homologs (Figs. S4, S6).

The recognition mode of the dinucleotide by ZF2 and ZF3 of CPSF-30 is distinct from those in other CCCH zinc fingers, such as that in Nab2 (35), the splicing factor muscleblind (MBNL1) (36, 37), and the AU-rich element binding protein TIS11d (38) (Fig. S7). While the position of the A1 base in ZF2 has approximate counterparts in the other structures, the position of A2 in CPSF-30 is unique. In fact, even the direction of the phosphate backbone is not the same among the bound RNAs. Overall, there appears to be substantial variability in ribonucleotide recognition by CCCH zinc fingers.

### CPSF-30 and WDR33 bind PAS RNA synergistically

Our structure shows that both CPSF-30 and WDR33 are directly involved in PAS recognition, while CPSF-160 is not in contact with this hexanucleotide. To test the importance of the three proteins in this binding, we carried out electrophoretic mobility shift assays (EMSAs) using the 17-mer RNA oligo, with a 3′-end fluorescent label to aid visualization. We used protein samples that were purified by gel filtration and showed homogeneous, non-aggregated behavior on the column (Fig. S8) for these assays. Our data showed that the CPSF-160–WDR33–CPSF-30 ternary complex has high affinity for the AAUAAA PAS RNA (Fig. 5A), while the affinity for the variant AAgAAA RNA is lower (Fig. 5B), consistent with earlier data (17). In contrast, the CPSF-160–WDR33 binary complex showed only weak binding (Fig. 5C), while the MBP-CPSF-30 fusion protein showed essentially no binding to the AAUAAA PAS RNA (Fig. 5D). A mixture of CPSF-160 and MBP-CPSF-30, which did not appear to form a binary complex (Fig. S8), also had very low affinity for the PAS RNA (Fig. 5E). In contrast, a mixture of the CPSF-160– WDR33 binary complex with a CPSF-30 deletion mutant (containing residues 33-170) that cannot form the ternary complex (Fig. 1B) showed appreciable binding for the PAS RNA (Fig. 5F), suggesting that the PAS RNA can stabilize the quaternary complex. Overall, the EMSA data indicate that the affinity of CPSF-30 and WDR33 alone or together with CPSF-160 for the PAS RNA is low, and the two proteins bind synergistically to the PAS RNA in the presence of CPSF-160.

**Figure 5.**
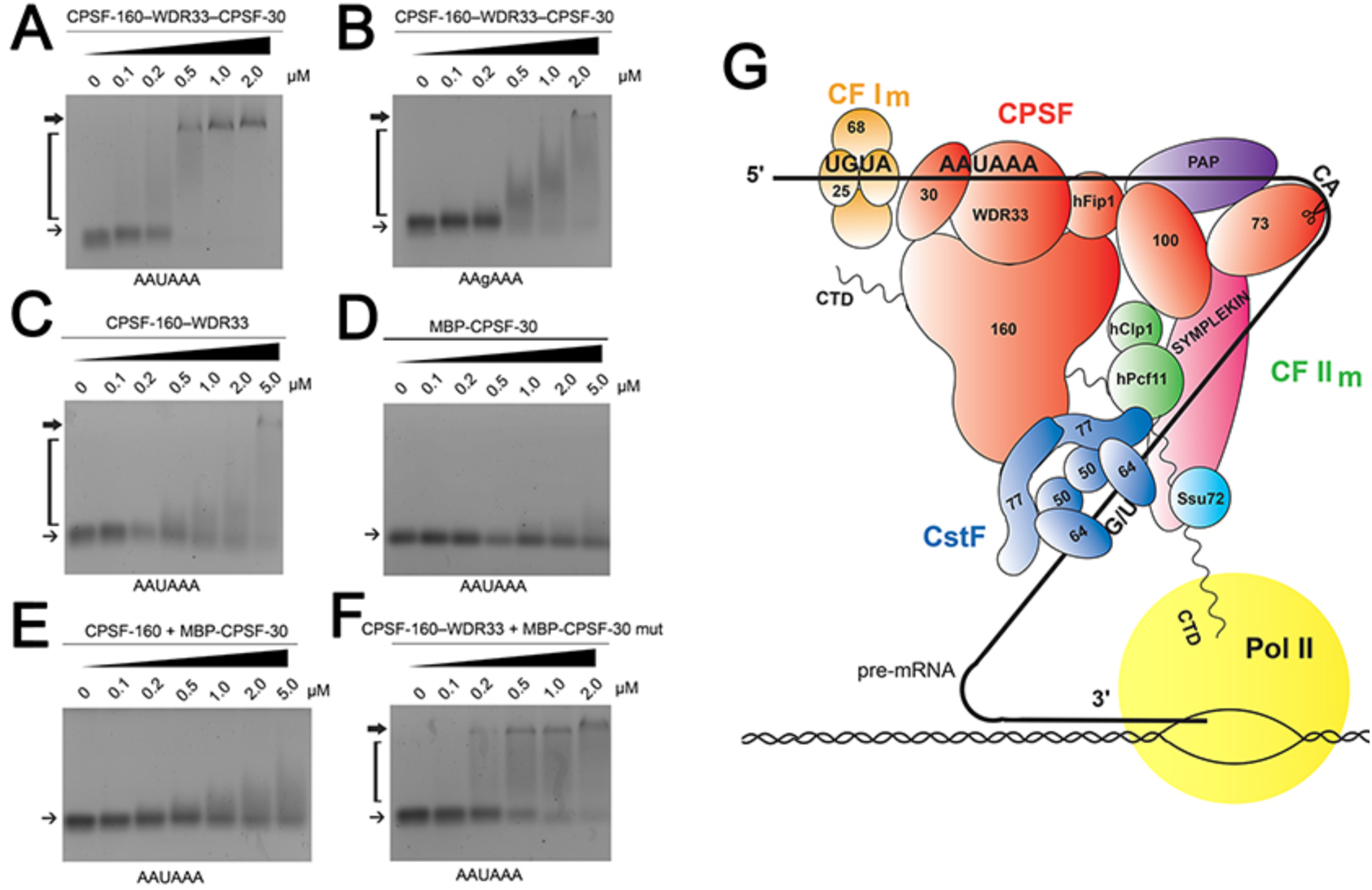
CPSF-30 and WDR33 bind the PAS RNA synergistically. **(A)**. Interaction between the CPSF-160–WDR33–CPSF-30 ternary complex and the 17-mer PAS RNA with a 3′fluorescent label by EMSA. The position of the RNA alone is indicated with the thin arrow, and that of the stable complex with the thick arrow. The bracket marks a smeared band, indicating complexes that dissociated during the electrophoresis. **(B)**. Interaction between the CPSF-160–WDR33–CPSF-30 ternary complex and a variant 17-mer PAS RNA, with AAgAAA hexamer. **(C)**. Interaction between the CPSF-160-WDR33 binary complex and the PAS RNA. **(D)**. Interaction between the MBP-CPSF-30 fusion protein and the PAS RNA. **(E)**. Interaction between a mixture of CPSF-160 and MBP-CPSF-30 and the PAS RNA. **(F)**. Interaction between a mixture of the CPSF-160–WDR33 binary complex and a mutant CPSF-30, MBP-CPSF-30(33-172), that cannot form the ternary complex and the PAS RNA. **(G)**. Schematic drawing of the mammalian canonical 3′-end processing machinery. The organization of the CPSF-160–WDR33–CPSF-30–PAS RNA complex is based on the structure described here.

### CPSF-160 is a scaffold

The overall structure of the CPSF-160–WDR33 complex has similarity to that of the DDB1–DDB2 complex that is important for DNA damage repair. The relative positions of BPA and BPC are similar to those in DDB1 (39) (Fig. S9). Large variations in the positioning of BPB are observed in DDB1, although we did not see significant flexibility in BPB of CPSF-160 in our studies here (Fig. 1C). DDB2 is bound between BPA and BPC of DDB1 as well, also using N-terminal extensions beyond the β-propeller domain. However, the position of the WD40 domain in WDR33 is different from that of DDB2 in the complex. Moreover, DDB2 binds DNA using its top face, and the positions occupied by CPSF-30 and PAS RNA in the quaternary complex has not been seen in the DDB1 complexes so far (Fig. S9). The structural similarity between CPSF-160–WDR33 and DDB1–DDB2 suggests that they might be related evolutionarily. A system used for DNA damage repair might have been re-purposed for RNA processing (or *vice versa*).

While CPSF-160 is not directly involved in interactions with the PAS, the structure indicates that it has an essential role in this recognition, as a scaffold to recruit WDR33 and CPSF-30 and position them correctly for binding the PAS RNA. Our EMSA data confirm that this pre-organization of the binding site by the CPSF-160 scaffold is crucial for high-affinity PAS recognition.

## Discussion

Our structure has illuminated the molecular mechanism for PAS recognition in pre-mRNA 3′-end processing, and it also provides a foundation for understanding and interpreting the biochemical data. Most importantly, earlier analyses have shown that AAUAAA is the predominant PAS among pre-mRNAs, with greater than 50% frequency (18, 19). The second most frequent PAS, AUUAAA, has a frequency of 16%, while all other PAS sequences have frequencies of less than 5%. Moreover, the U3 and A6 bases appear to be more conserved among the various PAS sequences. These observations are fully consistent with, and are explained by, our structure. For the AUUAAA hexamer, the recognition of the smaller U2 base could be mediated by a hydrogen-bond between its carbonyl group on C4 and the main-chain amide of Lys78 in CPSF-30, which would only require a small re-arrangement of the RNA.

The structure also shows that ZF2-ZF3 of CPSF-30 is crucial for PAS recognition. This is supported by earlier data showing that deletion of either zinc finger abolished cross-linking to RNA and polyadenylation activity (16). Moreover, our structure explains how influenza virus NS1A protein can disrupt host pre-mRNA 3′-end processing by sequestering ZF2-ZF3 of CPSF-30 (40, 41). The structure of ZF2-ZF3 in the quaternary complex here is similar to that in the complex with NS1A, while the bound position of NS1A clashes with the PAS RNA (Fig. 4H). Therefore, CPSF-30 can no longer participate in the recognition of the AAUAAA PAS in the presence of NS1A.

The WD40 domain of WDR33 contacts the U3-A6 base pair, although there is no direct recognition of the bases by hydrogen bonds. In comparison, the WD40 domain of Gemin5 was reported earlier to specifically recognize the Sm site in pre-snRNAs (42, 43), and the RNA is bound mostly at the interface of the tandem WD40 domains (44-46). These results suggest diverse modes for WD40 domains to interact with RNA, in addition to their roles in protein-protein and protein-DNA interactions.

The PAS is a relatively short motif with only six nucleotides. The structure shows that the motif is divided into three pairs of two nucleotides each, A1-A2, A4-A5, U3-A6, and each pair is recognized by a different protein component, which comes from two distinct proteins. Such a mode of interaction should enhance the specificity of the recognition, as each protein alone would have much lower affinity, which was demonstrated by our EMSA experiments. This binding mode also highlights the crucial importance of CPSF-160 as a scaffold, to pre-organize CPSF-30 and WDR33 and allow them to bind synergistically to AAUAAA with high-affinity. Therefore, the CPSF-160– WDR33–CPSF-30 ternary complex enables both high affinity and high selectivity in PAS recognition.

Many additional surface areas of CPSF-160 remain available in the complex, such as the bottom face of BPA, both faces of BPB and the sides of all three propellers (Fig. 1E), and CPSF-160 is likely to mediate interactions with other protein factors in the 3′-end processing machinery, for example CstF-77 (25, 47, 48). In fact, many of the equivalent surfaces in DDB1 have been found to interact with other proteins, and the protein bound between BPA and BPC does not have to contain a β-propeller either (30, 49) (Fig. S9). Besides recruiting other proteins, it remains to be seen whether CPSF-160 could also interact with other regions of the pre-mRNA near the PAS.

Overall, our studies have produced the first molecular insights into PAS recognition and the organization of the mPSF in the canonical 3′-end processing machinery (Fig. 5G). The hFip1 subunit of mPSF is not involved in PAS recognition although it may interact with the pre-mRNA near the PAS (50, 51). hFip1 also mediates the recruitment of the poly(A) polymerase (50, 52), enabling the polyadenylation reaction after cleavage. The mCF has strong interactions with the mPSF, forming CPSF (with symplekin), which together with CstF and other factors, defines the site of cleavage in the pre-mRNA. Further studies are needed to understand the molecular mechanism for these other steps in 3′-end processing.

## Methods

### Protein expression and purification

Human CPSF-160 and WDR33 (residues 1-572) were co-expressed in insect cells using Multibac technology (53) (Geneva Biotech). CPSF-160 and WDR33 were cloned into the pKL and pFL vectors, respectively, and a 6×His tag was added to the N terminus of WDR33. Bacmids for expressing CPSF-160 and WDR33 were generated in DH10EMBacY competent cells (Geneva Biotech) by transformation. Baculoviruses were generated by transfecting bacmids into Sf9 cells using Cellfectin II (Thermo Fisher Scientific). P1 viruses were cultured at 27 °C for 5 days, and P2 viruses for large-scale infection were amplified from P1 viruses in 50 ml Sf9 cells at 27 °C for 3 days. One liter of High5 cells (1.8×10^6^ cells ml^−1^) cultured in ESF 921 medium (Expression Systems) was infected with 15 ml CPSF160 P2 virus and 10 ml WDR33 P2 virus at 27 °C with constant shaking. Cells were harvested after 48 h by centrifugation at 2000 rpm for 13 min.

For purification, the cell pellet was re-suspended and lysed by sonication in 100 ml buffer containing 25 mM Tris (pH 7.9), 300 mM NaCl and one protease inhibitor cocktail tablet (Sigma). The cell lysate was then centrifuged at 13,000 rpm for 45 min at 4 °C. The supernatant was incubated with nickel beads for 1 h at 4 °C. The beads were then washed 4 times with 50 bed volumes of wash buffer (25 mM Tris (pH 7.9), 300 mM NaCl and 15 mM imidazole) and eluted with 25 mM Tris (pH 7.9), 150 mM NaCl and 250 mM imidazole. The protein was further purified by chromatography using a HiTrap Q column (GE Healthcare) and a Hiload 16/60 Superdex 200 column (GE Healthcare). The CPSF-160–WDR33 complex was concentrated to 4 mg ml^−1^ in buffer containing 25 mM Tris (pH 7.9), 300 mM NaCl and 5 mM DTT, and stored at −80 °C.

Human CPSF-30 (full length and residues 33-170) was cloned into the pRSFDuet vector (Novagen) and over-expressed in *E. coli* BL21 (DE3) Star cells. We used isoform 2 of CPSF-30, where residues 191-215 are absent. A 6×His tag followed by maltose binding protein (MBP) was added to the N terminus of CPSF-30, separated by a TEV protease cleavage site. Cell cultures were grown at 37 °C in LB-Agar (Sigma) containing 35 µg ml^−1^ kanamycin. When cell cultures reached an OD_600_ of 0.7~0.8, they were supplemented with 0.2 mM ZnCl_2_ for 15~30 min, and protein expression was then induced with 0.15 mM isopropyl β-D-1-thiogalactopyranoside (IPTG) at 18 °C overnight. Cells were harvested by centrifugation at 4000 rpm for 15 min. The MBP-CPSF-30 fusion protein was purified following a similar protocol as that used for the CPSF-160–WDR33 complex.

### CPSF-160–WDR33–CPSF-30–PAS RNA quaternary complex formation

Purified CPSF-160–WDR33 complex and MBP-CPSF-30 were mixed at a molar ratio of 1:3 in the presence or absence of the RNA oligonucleotide AACCUCCAAUAAACAAC (IDT). TEV protease was added at a ratio of 1:10 (w/w) with MBP-CPSF-30. The reaction mixture was incubated at room temperature for 2 h and then purified by gel filtration using a Superose 6 10/300 GL column (GE Healthcare), in a running buffer containing 25 mM Tris (pH 7.9), 350 mM NaCl and 5 mM DTT. Samples from the quaternary complex fractions were used for EM studies. The presence of RNA in the complex was confirmed with an A260/A280 ratio of 0.90, while the ratio was 0.64 if RNA was left out of the mixture.

### EM specimen preparation and data collection

The homogeneity of the samples was first examined by negative-stain EM with 0.7% (w/v) uranyl formate as described (54). A Philips CM10 electron microscope operated at 100 kV was used to collect 54 images for the CPSF-160-WDR33 sample. The PAS RNA was added to the sample before preparing the EM grids, but no RNA density was observed in the eventual cryo-EM 3D reconstruction. The images were recorded at a defocus of –1.5 µm on an XR16L-ActiveVu charge-coupled device camera (AMT) at a nominal magnification of 52,000x (calibrated pixel size of 2.42 Å on the specimen level) (Supplementary Fig. 1).

Before preparing grids for cryo-EM, the freshly purified protein samples were centrifuged at 13,000 g for 2 min to remove protein aggregates, and the protein concentration was measured with a NanoDrop spectrophotometer (Thermo Fisher Scientific). All specimens for cryo-EM were frozen with a Vitrobot Mark VI (FEI) set at 4 ºC and 100% humidity. Cryo-EM imaging was performed in the Cryo-EM Resource Center at the Rockefeller University using SerialEM (55) or, when stated, in the Simons Electron Microscopy Center at the New York Structural Biology Center using Leginon (56).

The CPSF-160–WDR33 sample was concentrated to 1.8 mg ml^−1^ and CHAPSO (8 mM final concentration) was added to prevent the particles from adopting preferred orientations in the ice layer (57). A 3-µl aliquot was applied to a glow-discharged Quantifoil 300 mesh 1.2/1.3 gold grid (Quantifoil). After 5 s, the grid was blotted for 3 s with a blot force setting of 0 and plunged into liquid ethane. Grids were screened with a Talos Arctica electron microscope, and 1,625 image stacks were collected on a 300-kV Titan Krios electron microscope (Thermo Fisher Scientific) at a defocus ranging from –1.3 to –2.7 µm. The images were recorded at a nominal magnification of 22,500x (calibrated pixel size of 1.3 Å on the specimen level) with a K2 Summit camera in super-resolution counting mode. Exposures of 10 s were dose-fractionated into 40 frames (250 ms per frame), with a dose rate of 8 electrons pixel^−1^ s^−1^ (~1.18 electrons Å^−2^ per frame), resulting in a total dose of 47 electrons Å^−2^.

For the CPSF-160–WDR33–CPSF-30–PAS RNA sample, a 4-µl aliquot at 0.18 mg ml^−1^ was applied to a glow-discharged Quantifoil 300 mesh 1.2/1.3 gold grid (Quantifoil). After 2 s, the grid was blotted for 4 s at a blot force setting of –2 and plunged into liquid ethane. Grids were screened with a Talos Arctica electron microscope, and 2,486 image stacks were collected on a Titan Krios electron microscope at the New York Structural Biology Center. The images were recorded with a K2 Summit camera in counting mode at a nominal magnification of 22,500x (calibrated pixel size of 1.07 Å on the specimen level) and a defocus range from –1.2 to –2.5 µm. Exposures of 10 s were dose-fractionated into 40 frames (250 ms per frame), with a dose rate of 8 electrons pixel^−1^ s^−1^ (~1.75 electrons Å^−2^ per frame), resulting in a total dose of 70 electrons Å^−2^.

### Image processing

EMAN2 (58) was used to pick 27,300 particles of the negatively stained CPSF-160–WDR33 complex. The particles were windowed into 80x80-pixel boxes, which were then resized to 64x64 pixels. After centering, the particles were subjected to classification with the iterative stable alignment and clustering (ISAC) algorithm (59), specifying 100 images per group and a pixel error threshold of 0.7. Eight generations produced 412 averages (Supplementary Figs. 1b). The class averages were used to calculate an initial model with the validation of individual parameter reproducibility (VIPER) algorithm implemented in SPARX (60) (Supplementary Fig. 1).

For the cryo-EM datasets, the image stacks were motion-corrected, dose-weighted, and binned over 2x2 pixels (binning was only performed for data collected in super-resolution mode) in MotionCor2 (61). The CTF parameters were determined with CTFFIND4 (62).

For the CPSF-160–WDR33 dataset collected with the Titan Krios, 372,707 particles were automatically picked with Gautomatch (http://www.mrc-lmb.cam.ac.uk/kzhang/Gautomatch/). After windowing the particles into 160x160-pixel boxes in RELION-2 (63), they were directly subjected to 3D classification into 6 classes, using the map obtained with the negative-stain EM dataset as initial model (2D classification in RELION-2 was performed but no particles were discarded). The orientation parameters of the particles in the largest class (containing 205,373 particles) were further refined, resulting in a final density map at 3.85 Å resolution according to the gold-standard Fourier shell correlation (FSC) curve and a cutoff of 0.143 (Supplementary Fig. 2). Particle polishing improved the resolution to 3.78 Å (Supplementary Fig. 3).

For the CPSF-160–WDR33–CPSF-30–PAS RNA dataset collected with the Titan Krios, 1,144,122 particles were automatically picked with Gautomatch using as templates 4 class averages that were obtained from 2D classification of the cryo-EM dataset of the CPSF-160–WDR33 complex. The particles were windowed into 192x192-pixel boxes and subjected to 2D classification in RELION-2. Particles in classes showing clear structural features were combined (529,190 particles) and subjected to 3D classification into 6 classes using as initial model the density map obtained with vitrified CPSF-160–WDR33 complex filtered to 30 Å resolution. Five classes produced maps with clear fine structure features, and the particles of these five classes were combined. Refinement yielded a density map at 3.36 Å resolution, which showed high-resolution features of the CPSF-160–WDR33 region. Two of the 6 classes showed additional density, likely representing CPSF-30 and PAS RNA. These two classes were combined and subjected to another round of 3D classification into 6 classes. Five of the resulting classes that showed the additional density were combined (173,632 particles) and refinement yielded a map at 3.42 Å resolution. To improve the density for the CPSF-30–PAS RNA region, additional refinement was performed using a soft mask containing BPC of CPSF-160, WDR33, CPSF-30 and PAS RNA. Particle polishing was performed but did not improve the map (Supplementary Fig. 2). The local resolution maps were calculated with the half maps in RELION-2.

### Model building and refinement

We used predicted structures for CPSF-160 from I-TASSER (64) and WDR33 from Phyre2 (65), and the crystal structure of ZF2-ZF3 of CPSF-30 (41) (PDB ID 2RHK) as starting models, and fitted them into the cryo-EM density map with Chimera (66) and Rosetta (67). All manual model building was performed with Coot (68). The atomic models were optimized by an iterative local rebuilding procedure in Rosetta using the map calculated with all the data and then further refined by using phenix.real_space_refine (69) against half-map 1 from RELION-2. FSC curves were calculated between the refined models and half map 1 (work), half map 2 (free), and the combined map (Supplementary Fig. 3). The statistics from the structure determination is summarized in Supplementary Table 1.

### Protein–RNA binding assays

Wild-type (AACCUCCAAUAAACAAC) and variant (AACCUCCAAgAAACAAC) 17-mer RNA oligos and the AAUAAA hexamer oligo (all with 3′-end 6-FAM label; IDT) were dissolved in DEPC-treated water. The gel shift assays were performed with 0.2 µM labeled wild-type or variant RNA and increasing concentrations of protein. The reactions were incubated at room temperature for 45 min in a 10 µl volume containing 20 mM Tris (pH 7.9), 200 mM NaCl, 1 mM MgCl_2_, 1 mM DTT, 5% (v/v) glycerol, and 0.1 mg ml^−1^ BSA. The samples were then supplemented with 1 µl 50% (v/v) glycerol and run on pre-chilled 0.6 % (w/v) TAE (Tris, acetate, EDTA) agarose gels at 140 V for 30 min. The gels were visualized on a Typhoon FLA 7000 (GE Healthcare).

### Sequence alignment

Alignment of selected sequences of CPSF-30 and WDR33 homologs was produced with Clustal Omega (70), presented with ESPript (71), and modified manually to include additional information.

## Acknowledgments

We thank Mark Ebrahim and J. Sotiris for help with data collection at the Evelyn Gruss Lipper Cryo-Electron Microscopy Resource Center at The Rockefeller University; Ed Eng, Bill Rice, Laura Kim and Bob Grassucci for help with data collection at the New York Structural Biology Center; Frank DiMaio for help with model building and refinement using Rosetta; Kehui Xiang for initial studies on CPSF-160. This research is supported by NIH grants R35GM118093 (to LT) and R01GM090056 (to Y. Shi). Some of this work was performed at the Simons Electron Microscopy Center and National Resource for Automated Molecular Microscopy located at the New York Structural Biology Center, supported by grants from the Simons Foundation (349247), NYSTAR, and the NIH National Institute of General Medical Sciences (GM103310) with additional support from the Agouron Institute (F00316) and NIH S10 OD019994.

